# High fat diet changes bacterial signatures in the murine pancreas

**DOI:** 10.1101/2022.04.29.490014

**Authors:** Torsten P.M. Scheithauer, Mark Davids, Maaike Winkelmeijer, Stefan Havik, Max Nieuwdorp, Hilde Herrema, Daniël H. van Raalte

**Affiliations:** Department of Experimental Vascular Medicine, Amsterdam University Medical Center (UMC), location Amsterdam Medical Center (AMC), Meibergdreef 9, 1105 AZ Amsterdam, The Netherlands; Department of Internal Medicine, Amsterdam University Medical Center (UMC), location Vrije University Medical Center (VUmc), The Netherlands

## Abstract

Hyperglycemia is caused by failure of pancreatic beta cells. Beta-cell inflammation contributes to beta-cell dysfunction, however (primary) immunogenic triggers are in large unknown. The gut microbiota is one potential source of pro-inflammatory molecules. A high-fat diet treatment increases pro-inflammatory bacteria in the gut microbiome, parts of which could migrate to the pancreatic beta-cell. In the present study, the bacterial DNA signature in the pancreas and intestine of C57BL6/J mice was analyzed. Mice were fed a high-fat diet (60% kcal fat) or a regular chow diet for 12 weeks. We took several precautions to avoid and map contamination. The gut microbiota was affected by high-fat diet as following: We observed several common intestinal ASVs in the pancreatic tissue. Although the pancreatic ASVs do not correlate exactly with the gut ASVs, our data implicate that the pancreas contains bacterial DNA and that this signature is altered in high-fat diet fed mice (PERMONOVA, p = 0.037; betadisper, p=0.029). Gut derived bacterial DNA might end up in the pancreas at some point in time. Hence, this work supports the concept of translocation of bacterial DNA to the pancreas, which might contribute to inflammation and dysfunction of pancreatic beta-cells.

## Main text

Obesity^1^ and Type 2 diabetes (T2D)^2^ are global pandemics that are challenging health systems worldwide. The obesity-driven insulin resistance increases insulin demand from pancreatic beta cells^3^. Further, inflammation induces beta-cell dysfunction and prevents increment in insulin secretion necessary to overcome insulin resistance^4^. Underlying causes for beta-cell inflammation are mostly unknown. Hyperglycemia, saturated fatty acids and islet amyloid polypeptide are discussed as pro-inflammatory signals^5^.

Recent findings suggest that the gut microbiota might be involved in regulation of host glucose metabolism. Numerous studies have reported a relation between so called “gut dysbiosis” and the development of T2D^6,7^. Individuals with obesity and T2D have lower microbial diversity, while showing increased abundance of potentially pathogenic Gram-negative bacteria^8^. Bacterial translocation from the intestine into metabolic active tissues might lead to inflammation and in turn function loss^9-11^. However, measuring bacterial DNA in eukaryotic tissue is still very controversial amongst others due to technical limitations.

The pancreas was thought to be sterile, but recent studies find bacterial signatures^12-15^. Gao *et al* (2022) measured extracellular vesicles with microbial DNA in human and murine pancreatic islets^16^. Further, tumours harvested from pancreatic cancer patients display a diverse bacteriome^12,13^ and mycobiome^14^ that is associated with disease activity^15^. In light of extensive efforts in our own group on the role of the gut microbiome in regulation glucose and insulin regulation in mice and humans^6,17-19^, we here designed a controlled experiment to address pancreas bacterial abundance in mice.

We used male six-week-old C57BL6J mice, which were either fed a normal chow diet or a high-fat diet (HFD, 60% kcal fat) for 12 weeks. Mice were housed with two mice per cage to reduce cage effects facilitated by coprophagy. During termination of the mice, sterile and dedicated instruments were used per organ. Instruments were thoroughly cleaned between mice using a protocol that included heating, bleaching and rinsing with DNA free water. Lysis buffers, cleaning reagents and fur were included in the sequencing run to control for contaminants.

As expected, HFD fed mice had significantly higher body weights (**Figure 1A**) and higher blood glucose levels compared to lean mice (**Figure 1B and 1C**). Both, duodenal and caecal microbiome were different between lean and obese mice (**Figure 1D**). Composition and dispersion were different when comparing the pancreas of obese and lean mice (**Figure 1E and 1F**, PERMONOVA, p = 0.037; betadisper, p=0.029). Most of the pancreatic signature could be traced to reagent contaminants and on average 80% of the reads were discarded. No obvious contaminating source for the remaining bacterial signature could be identified. Pancreatic ASVs represented bacterial species typically present in the gut and differed between chow and HFD (**Figure 1F**). We therefore conclude that the pancreas contains gut bacterial DNA. This could not be explained by technical confounders like amplicon yields or contamination rates (**Figure 1F**), suggesting that a HFD changes translocating bacterial DNA.

**Figure 1:**
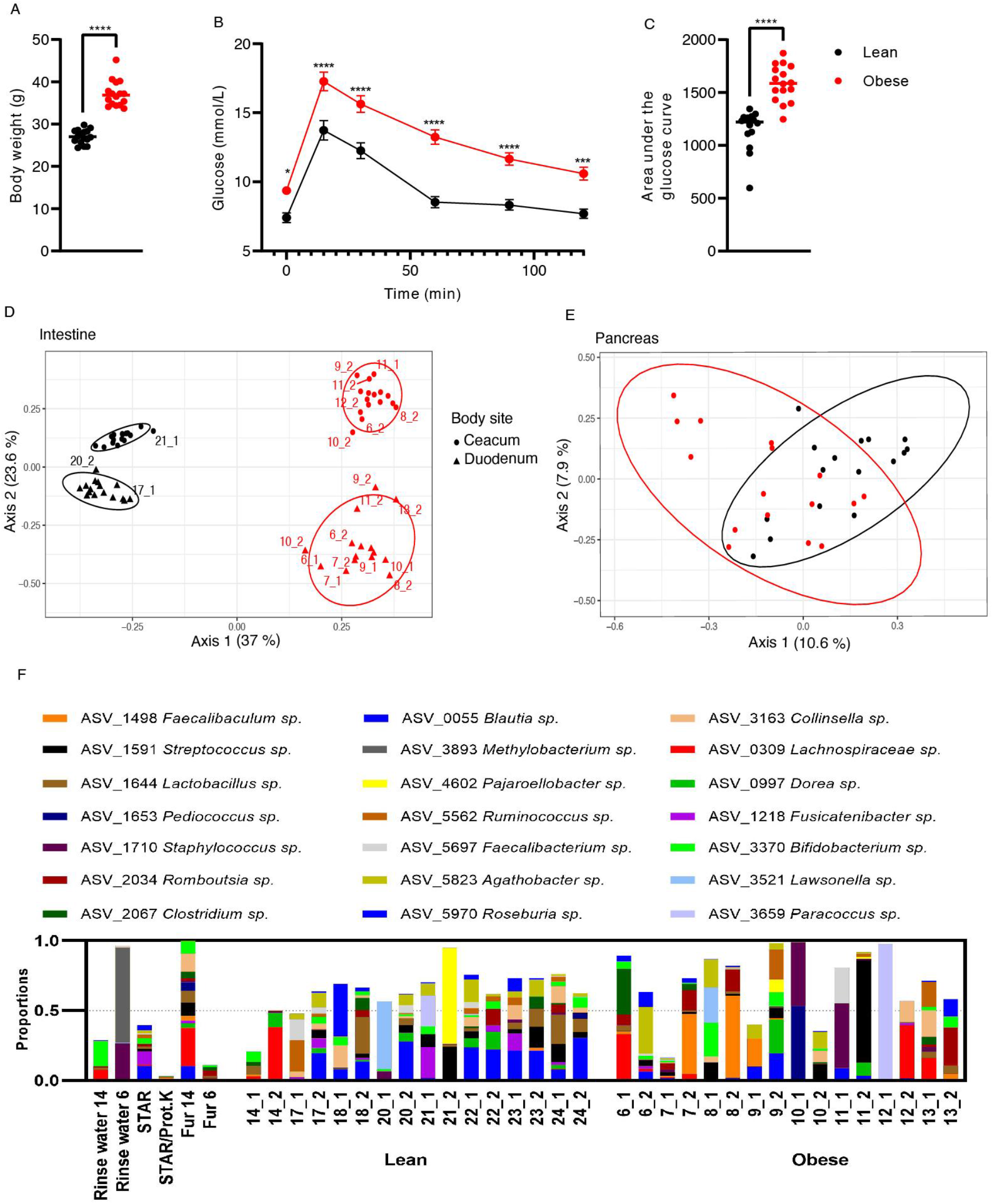
High fat diet changes the bacterial signature of the murine pancreas. C57BL6J mice were fed a 60% kcal high fat diet or a normal chow diet for 12 weeks (n = 16). Mice were dissected aseptically. 16s rRNA was sequenced using the Illumina MiSeq platform. (A) Body weight was higher in obese mice. (B, C) Obese mice were more glucose intolerant than lean mice. (D) The duodenal and caecal microbiome differed between groups as shown by 16S rRNA miSeq sequencing. Principal Coordinates Analysis (PCoA) analysis shows different composition between the two tissues and groups. (E) The bacterial signature in the pancreas is different between the two groups. (F) The bacterial signature in the pancreas as well as control samples were sequenced (rinse water for cleaning the instruments, STAR lysis buffer, proteinase K, fur of the mice). Individual mice are labelled with the cage number, followed by the mouse. Mice were housed in pairs.

Our results support the notion that bacteria components might translocate into the pancreas. However, technical challenges have to be noted. Although we included several controls, we cannot exclude that contaminants entered during the sample handling protocols^20^. Bacterial signatures have been also noted in the placenta^20^. Although these signatures could in large be traced back to contamination, it could not be fully excluded that bacterial DNA might be present in this sterile assumed tissue^20^. Of note, in our work, the ASVs in the pancreas did not exactly match the ASVs in the duodenum and caecum (data not shown). We speculate that pancreatic ASVs represent translocation events that occurred prior to termination of the mice. This hypothesis would need dedicated tracing experiments.

Potential routes of translocation include the duct between duodenum and pancreas as well as reduced intestinal tight junction expression in obese settings^21^. A recent paper suggests that microbial DNA-containing extracellular vesicles translocate into the pancreatic islets, where they induce inflammation and beta-cell dysfunction^16^. This effect is mediated via macrophages. It is plausible that (activated) immune cells transport bacterial parts from other tissues like the intestine^22^ into pancreatic tissue. Although not tested yet, increased pathogenicity of the gut microbiota might lead to increased mobility of gut bacteria and in turn bacterial presence in the pancreas. However, current techniques might not be sensitive enough to detect small amounts of bacterial DNA in large portions of eukaryotic DNA. Lastly, a HFD also increases the fat content of the pancreas. It is not known whether tissue composition also has an influence on the sequencing process.

In conclusion, our data imply that the murine pancreas contains gut bacterial DNA and that this bacterial signature differs in HFD-fed mice compared to regular chow fed mice. Although this concept needs further (technical) validation, our findings coincide with recent observations^16^ and open up the possibility that bacteria might contribute to inflammation and dysfunction of pancreatic beta-cells.^23,24^.

## Acknowledgments

DHvR was supported by a fellowship of the Dutch Diabetes Foundation (2015.81) and EU Marie Curie Program (H2020-MSCA-IF-2015). HH is supported by a Senior Fellowship of the Dutch Diabetes Research Foundation (2019.82.004). MN is supported by a ZONMW VICI grant 2020 [09150182010020].

## Methods

### Animals

C57BL/6J mice were purchased from Charles River (France) and maintained under specific pathogen free conditions in the S-building of the Amsterdam UMC, location AMC. All animals were housed in pairs, under a 12h light/dark cycle. All mice were fed a normal chow diet until week six, afterwards mice were kept on a normal chow diet or put on a 60% kcal high fat diet (ResearchDiets, USA) for 12 weeks. Only male mice were included in this study. Animal work was performed in accordance with the Central Commission for Animal Experiments (CCD, The Netherlands). Animals were fasted for 4h to measure fasting glucose. A intraperitoneal glucose tolerance (IPGTT) test was performed in the last week (1.5 g/kg glucose in saline). Mice were sacrificed in the laminar flow hood. Instruments were extensively cleaned by heating to 240-270°C and bleach for several minutes, followed by rinsing with ultrapure water (ThermoFisher, USA). Different instruments were used for opening up the mice, extracting the pancreas and the intestine. Instruments were cleaned between every mouse. Organs were snap frozen in sterile tubes in liquid nitrogen and stored until further use.

### 16S rRNA gene amplicon sequencing

DNA was extracted from 50-100 mg tissue using a repeated bead beating protocol (method 5)^25^. DNA was also isolated from rinse water of instruments, the fur of the mice and the lysis buffer to control from other contamination sources. DNA was purified using Maxwell RSC Whole Blood DNA Kit. Bacterial profiles were obtained by sequencing the V34 region of the 16S rRNA gene, obtained by PCR using the 515F and 806R primers designed for dual indexing^26^, on an Illumina MiSeq instrument (Illumina RTA v1.17.28; MCS v2.5) with the V3 Illumina kit (2×251 bp paired-end reads). The PCR was performed in a total volume of 30 μl containing 1× High Fidelity buffer (Thermo Fisher Scientific, Waltham, MA, USA); 1 μl deoxynucleoside triphosphate (dNTP) mix (10 mM; Promega, Leiden, The Netherlands); 1 U of Phusion green high-fidelity DNA polymerase (Thermo Fisher Scientific, Waltham, MA, USA); 500 nM the forward 8-nucleotide (nt) sample-specific barcode primer containing the Illumina adapter, pad, and link (341F [5′-CCTACGGGNGGCWGCAG-3′]); 500 nM the reverse 8-nt sample-specific barcode primer containing the Illumina adapter, pad, and link (805R [5′-GACTACHVGGGTATCTAATCC-3′]); 100 ng/μl of template DNA; and nuclease-free water. The amplification program was as follows: an initial denaturation step at 98°C for 30 s; 30 cycles of denaturation at 98°C for 10 s, annealing at 55°C for 20 s, and elongation at 72°C for 90 s; and an extension step at 72°C for 10 min.

### Bioinformatic processing and statistical analysis

Forward and reverse reads were truncated to 240 and 210 bases, respectively and merged using USEARCH<u>^18^</u>. Merged reads that did not pass the Illumina chastity filter, had an expected error rate higher than 2, or were shorter than 380 bases were filtered. Amplified Sequence Variants (ASVs) were inferred for each sample individually with a minimum abundance of four reads. Unfiltered reads were then mapped against the collective ASV set to determine abundances. Taxonomy was assigned using the RDP classifier^<u>19</u>^ and SILVA^<u>20</u>^ 16S ribosomal database V132. The AVS table, taxonomy and sample data were integrated using the ‘phyloseq’ R package (v.1.34.0) for further downstream curation and statistical analysis. ASV sequences previously identified, using decontam ^27^, as contaminants in the reagents, together with sequences from common reagent contaminants of the genera Ralstonia, Bradyrhizobium and Pelemonas, were marked as contaminants and removed from the ASV table. Differences in composition and dispersion were tested using vegans (V2.5.7) adonis and betadisp function.

